# Characterizing biofilm interactions between *Ralstonia insidiosa* and *Chryseobacterium gleum*

**DOI:** 10.1101/2022.10.11.511742

**Authors:** Andrea Foote, Kristin Schutz, Zirui Zhao, Pauline DiGianivittorio, Bethany R. Korwin-Mihavics, John J. LiPuma, Matthew J. Wargo

**Affiliations:** Cellular, Molecular, and Biomedical Sciences Graduate Program, University of Vermont Larner College of Medicine, Burlington, Vermont, USA; Department of Microbiology and Molecular Genetics, University of Vermont Larner College of Medicine, Burlington, Vermont, USA; Department of Biology, University of Vermont, Burlington, Vermont, USA; Department of Pediatrics, University of Michigan Medical School, Ann Arbor, Michigan, USA

**Keywords:** tap water microbiology, bacterial ecology, dual-species biofilms, bacterial interactions

## Abstract

*Ralstonia insidiosa* and *Chryseobacterium gleum* are bacterial species commonly found in potable water systems and these two species contribute to the robustness of biofilm formation in a model six-species community from the International Space Station (ISS) potable water system. Here, we set about characterizing the interaction between these two ISS-derived strains and examining the extent to which this interaction extends to other strains and species in these two genera. The enhanced biofilm formation between the ISS strains of *R. insidiosa* and *C. gleum* is robust to starting inoculum and temperature, occurs in some but not all tested growth media, and evidence does not support a soluble mediator or co-aggregation mechanism. These findings shed light on the ISS *R. insidiosa* and *C. gleum* interaction, though such enhancement is not common between these species based on our examination of other *R. insidiosa* and *C. gleum* strains, as well as other species of Ralstonia and Chryseobacterium. Thus, while the findings presented here increase our understanding of the ISS potable water model system, not all our findings are broadly extrapolatable to strains found outside of the ISS.

**Importance:** Biofilms present in drinking water systems and terminal fixtures are important for human health, pipe corrosion, and water taste. Here we examine the enhanced biofilm of cu-cultures for two very common bacteria from potable water systems, *Ralstonia insidiosa* and *Chryseobacterium gleum*. While strains originally isolated on the International Space Station show enhanced dual-species biofilm formation, terrestrial strains do not show the same interaction properties. This study contributes to our understanding of these two species in both dual and mono-culture biofilm formation.

## Introduction

Multispecies, surface-attached biofilms are an important part of the built environment, particularly the potable water system that includes the municipal delivery system, building plumbing, terminal fixtures, appliances, and associated surfaces [1–6]. Potable water system biofilms contribute to alterations in material corrosion, water properties, and health of those drinking and bathing in the water [7]. While the microbial communities within these systems are diverse, there are a number of taxa that are common across wide swaths of geography and water chemistry, including the genera *Ralstonia* and *Chryseobacterium* [8]. *Ralstonia insidiosa* and its close relative, *Ralstonia pickettii*, are beta proteobacteria that are common in water systems and other parts of the built environment [9, 10] and can be found infrequently as opportunistic pathogens in human infections [11–14]. *Chryseobacterium gleum* (formerly, *Flavobacterium gleum*) in the phylum Bacteroidetes, is present in similar environments as *R. insidiosa* and can also be present in human infections [15–17], though less frequently.

*R. insidiosa* promotes biofilms in conjunction with a number of species including *Listeria monocytogenes, Salmonella enterica*, and *Escherichia coli* [18–22]. In most of these cases, *R. insidiosa* has been reported as a physical bridge between cells of the other species with components of the biofilm matrix predicted as driving the enhanced biofilms [21, 22]. *Chryseobacterium* species are reported as strong biofilm formers in single-species cultures [17], but their enhancement of multispecies biofilms has not been reported as frequently [23].

We previously reported that strains of *R. insidiosa* isolate and *C. gleum* from the International Space Station (ISS) were capable of enhanced dual-species biofilm formation, though neither was capable of robust biofilm formation alone [24]. This interaction is a critical facet for robustness of biofilm formation in this six-species model drinking water community [24]. Here we characterize the interaction between *R. insidiosa* and *C. gleum* in relation to biofilm formation, examining potential broad mechanisms, the dependence of the interaction on growth conditions, and the broader applicability of this interaction by testing other strains and species within these two genera.

## Methods

### Bacterial strains and maintenance conditions

*R. insidiosa* 130770013-1 was isolated from the ISS Potable Water Delivery System and *C. gleum* 113330055-2 was isolated from the Russian SVO-ZV module on the ISS, and both identified by the Microbiology Laboratory at the NASA Johnson Space Center (Houston, TX) [1]. The strain numbers listed after each species are for the strain database at the Johnson Space Center, and strains may be requested directly using these designations. Terrestrial strains of *R. insidiosa, R. pickettii, C. gleum, Chryseobacterium indologenes, Chryseobacterium meningosepticum*, and *Ralstonia* strains that cannot be identified as belonging to one of the currently named species in this genus were acquired from the Cystic Fibrosis Foundation *Burkholderia cepacia* Research Laboratory and Repository at the University of Michigan. Details on all strains presented in **Table 1**. All bacteria were stored in 20% glycerol stocks at −80°C and were recovered on R2A plates at 30°C.

**Table 1:**
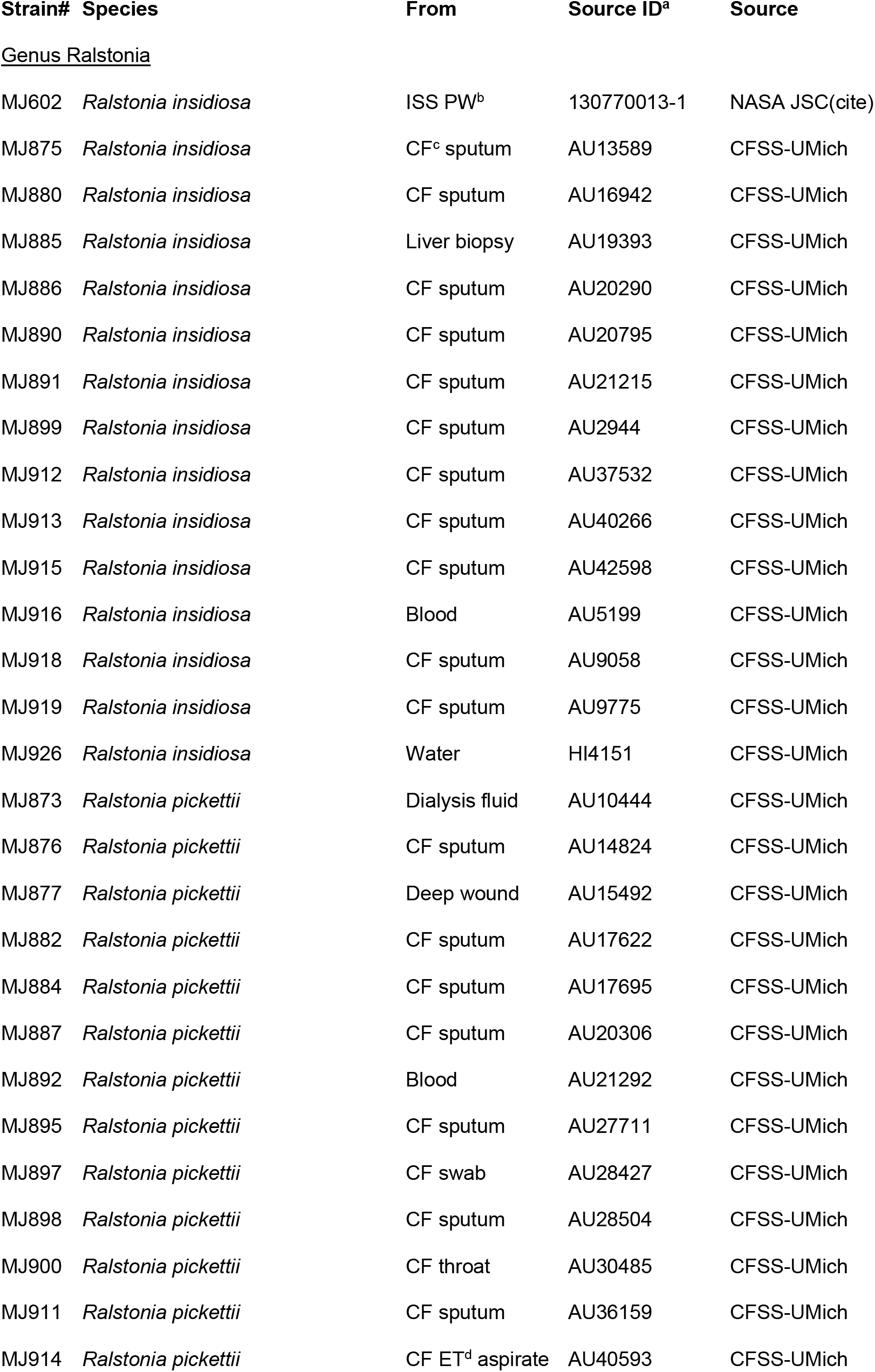
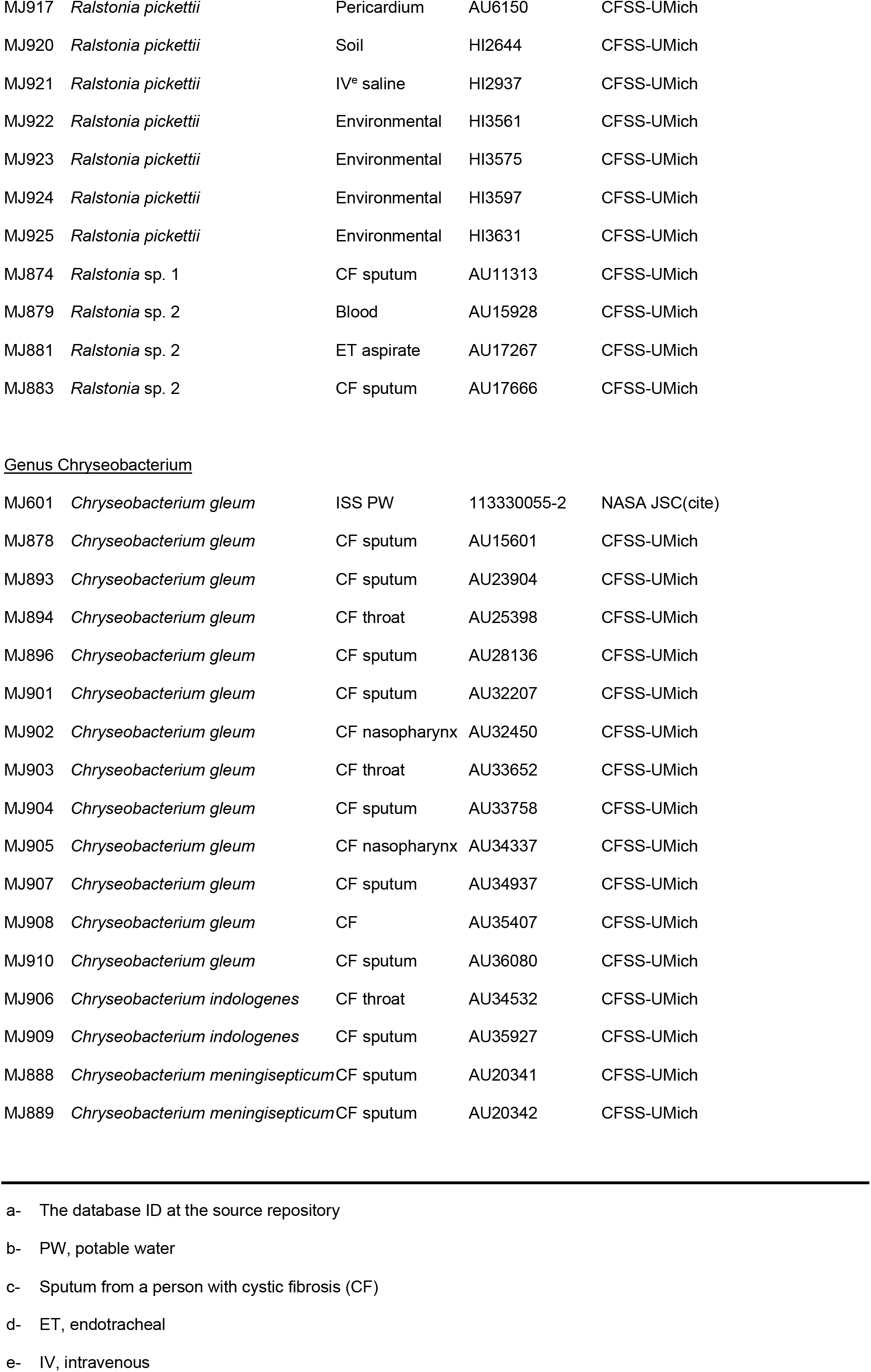
Description of strains used in this study

For all experiments below, the R2A plates were used as a source of inoculation into MRLP media [24] (2/3^rd^ modified MOPS media [25], 1/3^rd^ R2B, with 2% LB and 10 mM sodium pyruvate; in composition, this medium comprises 26.4 mM MOPS, 2.64 mM tricine, 6.28 mM NH_4_Cl, 34.7 mM NaCl, 0.35 mM MgCl_2_, 0.19 mM K_2_SO_4_, 1.57 mM K_2_HPO_4_, 10.91 mM sodium pyruvate, 0.94 mM glucose, 32 μM MgSO_4_, 21.1 μM CaCl_2_, 6 μM FeSO_4_, 5.28 μM FeCl_2_, 0.33 g l^-1^ peptone, 0.27 g l^-1^ yeast extract, 0.20 g l^-1^ tryptone, 0.17 g l^-1^ soluble starch, and 0.66X MOPS medium micronutrients) and cells were grown overnight in 3 ml culture volume at 30°C with orbital shaking of angled, loosely capped 18 x 150 mm glass tubes. Some experiments were also conducted in a high calcium version of MRLP where the final CaCl_2_ concentration was 400 μM. The higher calcium concentration increases monoculture biofilm formation for *C. gleum* as described in the results section, but co-culture enhancement is still apparent. Calcium concentration is noted in each figure legend.

Other media formulations were also tested to assess the range of conditions within which coculture biofilm enhancement could be seen. These included versions of MRLP where the 2/3 volume of minimal media was switched from MOPS to M9 or M63 to generate M9-RLP and M63-RLP, respectively. Additionally, RL media was used which was 1/3 R2B with 2% LB with the remainder of the volume made up from distilled-deionized water.

### Biofilm initiation, quantification, and cell numbers

For 96-well format static biofilms, overnight MRLP-grown cultures were used as the starting source for biofilm experiments. The optical density of each species was determined and normalized to generate an initial OD_600_ of 0.05 for each species member in MRLP. After pipetting 150 μl of the species mixtures into flexible 96-well plates, the plates were placed into humidified chambers and incubated under static conditions (no shaking) at 30 °C for 24 hours. To measure biofilm formation, the crystal violet staining protocol was used as described by O’Toole and colleagues [26]. Biofilms were quantified by dissolving the crystal violet with 10% acetic acid and measuring absorbance at 550 nm.

For rotating wall vessel (RWV) biofilms, cells were grown overnight as for the 96-well biofilms and then inoculated into 10 ml RWVs (Synthecon Inc., ethanol-sterilized and dried) at a starting inoculum of 0.008 OD_600_ per species. RWVs were rotated in the vertical orientation at room temperature overnight. In conducting the RWV experiments, we observed that most of the biomass, particularly in the dual-species culture, was present as a biofilm on the air-permeable membrane of the vessel. To measure these biofilms, the membrane was rinsed with sterile media and then scraped with a silicon cell scraper into a Petri dish containing additional sterile media. The contents of the Petri dish were then collected by centrifugation and pellets stained with crystal violet as above. After staining, pellets were washed twice before dissolution of the crystal violet in 10% acetic acid. Due to the large quantity of biofilm material, the crystal violet was quantified by two-fold serial dilutions, which were then corrected to total absorbance by multiplication with the respective serial dilution factor.

For biofilms on glass coverslips, 22 mm square coverslips were sterilized in ethanol and dried prior to insertion into 12-well plates. The coverslips fit snugly in the wells but rest slightly off of vertical. Each well contained 2.2 ml of high calcium MRLP, which was enough to reach about halfway up the coverslip. After inoculation as described for the 96-well biofilms, the 12-well dished were incubated statically at 30°C in a humidified chamber for 20-24 hours.

To determine total cell number in the static 96-well plates (planktonic + biofilm), cells were gathered by resuspending the contents of each well via vigorous pipetting and scraping the well walls with a pipette tip. As measured by crystal violet staining, < 1% of the biomass remained after this treatment. The scraped well contents were vortexed to resuspend and serially diluted in R2B before plating onto R2A. For wells with both species, CFU counts for each species were determined by the obvious colony color difference (orange-yellow for *C. gleum*, and white/beige for *R. insidiosa*).

### Statistical analysis and data visualization

All data visualization and statistical analyses were conducted in GraphPad Prism v9 using one-way or two-way ANOVA with Tukey, Sidak, Bonferroni, or Dunnett’s post-testing as described in individual figures. Values of p below 0.05 were considered statistically significant. Data is presented, when possible, with bars representing the mean and either biological replicates or all experimental replicates shown as individual data point overlain. Even when all experimental replicates are depicted, statistical analysis is conducted using the means of each biological replicate only. Where individual data points are not shown for clarity and figure size considerations, error bars represent standard deviation.

### Generation of mScarlet-expressing *R. insidiosa*

We used *attTn7-based* chromosomal integration to generate stable fluorescent *R. insidiosa* using the general method for Gram-negative bacteria [27]. Briefly, we mixed *R. insidiosa* with *E. coli* S17-1 carrying pMRE-Tn7-145 [27] and allowed conjugation to occur on an LB plate with 0.1% L-arabinose overnight. The spot was scraped, resuspended, and *R. insidiosa* carrying the *attTn7* insertion were selected by plating on MOPS agar with 25 mM pyruvate, 10 mM glucose, 25 μg/ml gentamicin, and 10 μg/ml chloramphenicol. Colonies were picked and screened for presence of the *R. insidiosa* specific PCR product [11], the integration cassette via PCR for the gentamicin resistance gene (GmR-check-F 5’-TCTTCCCGTATGCCCAACTT-3’ and GmR-check-R 5’-ACCTACTCCCAACATCAGCC-3’), and fluorescence concordant with mScarlet (using 565 excitation and 600 emission using a Biotek H1 multimode plate reader).

### Microscopy

Fluorescence microscopy was carried out on a Nikon A1R-ER confocal laser microscope housed in the UVM Microscopy Imaging Center using the RFP laser and emission filter for mScarlet which, while not optimized for mScarlet, still allowed visualization of the mScarlet signal. Images were captured and analyzed with Nikon NIS-Elements software.

## Results

### Co-culture of the ISS *Chryseobacterium gleum* and *Ralstonia insidiosa* leads to enhanced biofilm formation

We had previously observed that the ISS isolates of *R. insidiosa* and *C. gleum* in our six-member potable water biofilm community demonstrated enhanced biofilm formation together and very little when grown alone [24]. The goal of this project was to extend the characterization of this interaction. Partly to aid in this endeavor, we generated a clone of the ISS *R. insidiosa* carrying mScarlet at the *attTn7* site. Both the parent strain of *R. insidiosa* and the *R. insidiosa* with mScarlet showed enhanced biofilm formation with *C. gleum* in flexible 96-well dish biofilms (**Figure 1A**). This enhanced biofilm formation is not due solely to growth stimulation of either species during co-culture (**Figure 1B**).

**Figure 1.**
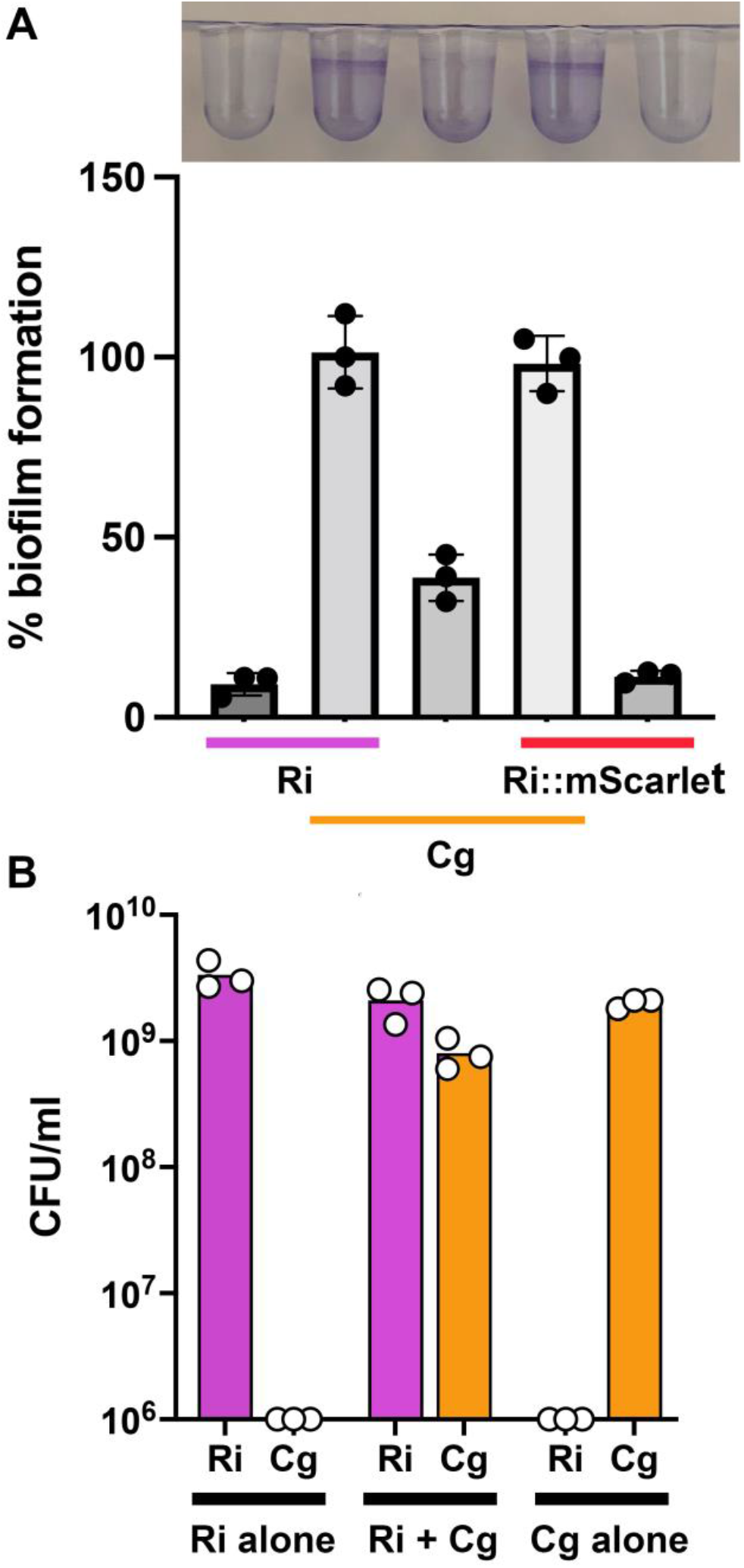
Enhanced biofilm formation during *Ralstonia insidiosa* and *Chryseobacterium gleum* coculture. The ISS isolates of *R. insidiosa* (Ri) and *C. gleum* (Cg) grown alone or cocultured in high-calcium MRLP medium for 24 hours. (**A**) (Top) Representative crystal violet stain of these static biofilms grown in flexible 96-well dishes, with samples in the same order as the graph. (Bottom) Quantification of the crystal violet biofilm absorbance normalized to *R. insidiosa+C. gleum* as 100% biofilm formation. Integration of *mScarlet* at the *attTn7* site (Ri::mScarlet) does not impact *R. insidiosa’s* biofilm formation or interaction with *C. gleum*. **(B)** Quantification of planktonic and biofilm cells from the mono- or co-culture wells. After scraping, vortexing, and resuspension, colony forming units were counted by serial dilution plating and subsequent counting. The y-axis starts at 10^6^ CFU as that was the detection limit for these dilutions. For each panel, each individual dot is the mean of three technical replicates on a separate experimental day. For (A), using ANOVA with a Sidak’s post-test, the biofilm formed by the co-cultures was significantly higher than that formed by the respective monocultures (p < 0.001) and was also significantly higher than the sum of the monoculture biofilms (p < 0.005). For (B), using a two-way ANOVA with Tukey’s post-test, the co-culture does not significantly impact the CFU of the individual bacteria compared to monoculture and the total cells per well are also not significantly altered.

To examine the enhanced biofilm microscopically, we grew each biofilm on glass coverslips positioned roughly vertically in 12-well dishes. To visually assess biofilm formation, we stained duplicate coverslips with crystal violet. Enhanced biofilm is apparent in the co-cultures, and we also note that *R. insidiosa attTn7::mScarlet* forms a stronger mono-culture biofilm on glass than it does on plastic (**Figure 2A**), which is not different than the untagged *R. insidiosa* on glass. The doublet line, most apparent in the co-culture, is formed because the coverslip rests at an angle off vertical; the surface film intercepts at slightly different positions on each side of the coverslip. For fluorescent microscopy, we wiped one side thoroughly with paper cloth after fixation to remove material from the side not being viewed. Imaging of the co-culture biofilm showed red-fluorescent cells that were co-stained with DAPI (*R. insidiosa*), as well as cells fluorescing solely with DAPI (*C. gleum*) (**Figure 2B**). Like other examples of *R. insidiosa* enhancing biofilm formation [19–22], this dual-species biofilm is intermixed.

**Figure 2.**
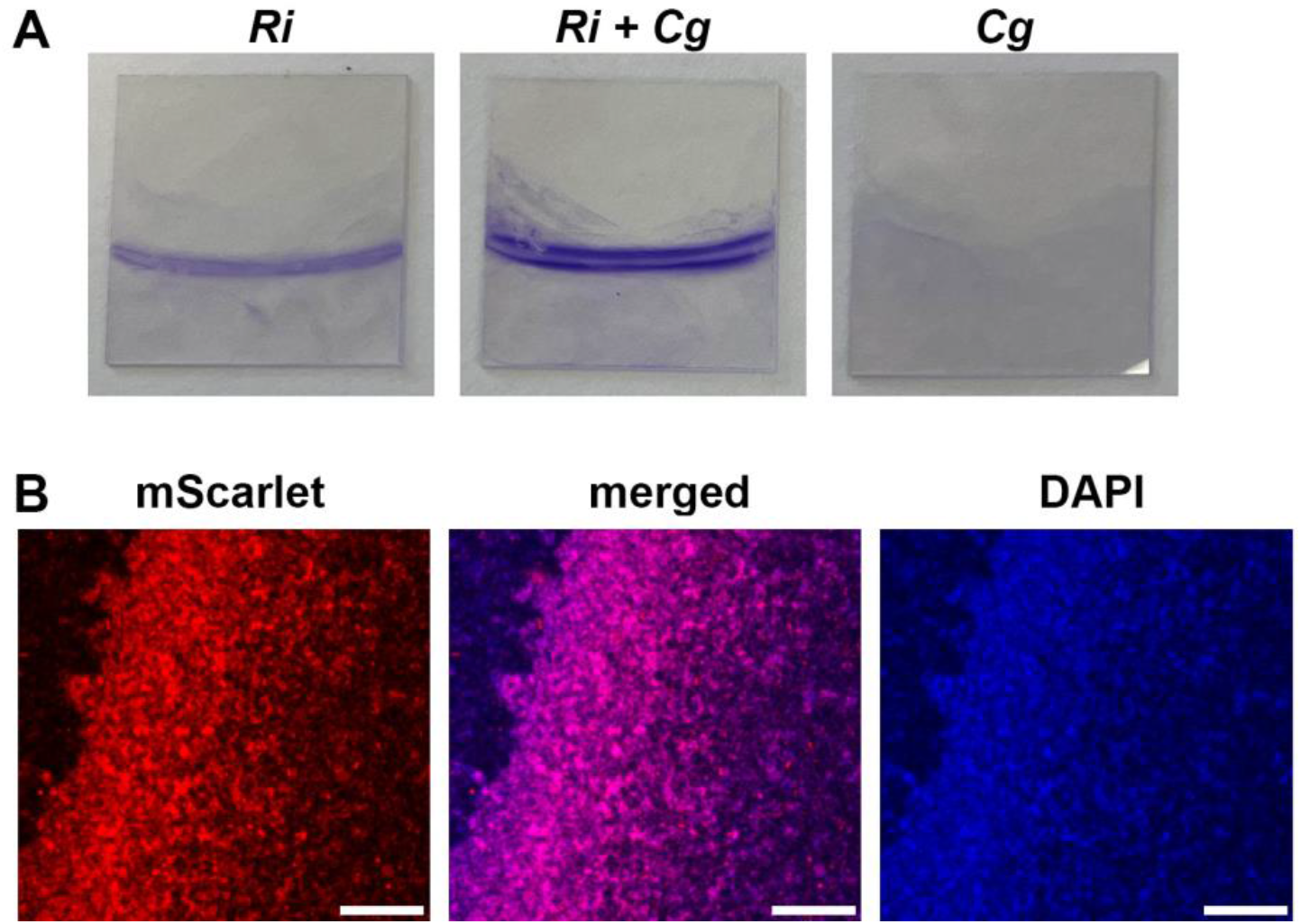
Enhanced dual-species biofilm formation on glass coverslips by *C. gleum* and *R. insidiosa attTn7::mScarlet*. The ISS isolates *C. gleum* (Cg) and the mScarlet-expressing *R. insidiosa (Ri*) were grown alone or cocultured in high-calcium MRLP medium for 24 hours in 12-well plates with a vertically oriented coverslip in each well. **(A)** Crystal violet staining of the coverslip biofilm. Note that *R. insidiosa* forms more biofilm on glass than on plastic (Figure 1). **(B)** One example Z-section of the dual-species biofilm taken approximately 3 μm above the coverslip using confocal microscopy. The air-liquid interface runs roughly diagonally on a line rotated slightly clockwise from vertical with the air side on the left. Scale bar = 100 μm.

### The impact of calcium concentration on *C. gleum* biofilm formation

During compilation of the data and final sets of experimentation, it was noticed that while enhanced biofilm was always obvious between these two strains of bacteria, the baseline biofilm formation by *C. gleum* was very different between different experimenters and compared to our previous work [24]. A deep dive through lab notebooks uncovered the likely difference as two different recipes for our MRLP media in the lab that differed only in their final calcium concentrations: 21.1 μM vs 400 μM. To test whether this calcium difference was able to explain the *C. gleum* biofilm differences between experimenters, we tested freshly made versions of both MOPS recipes. High calcium increases *C. gleum* biofilm formation (**Figure 3**). There is roughly 5-fold higher biofilm formation by *C. gleum* grown in high calcium versus in low calcium. The dual-species biofilms were also enhanced by high calcium growth, but biofilm for *R. insidiosa* remained unchanged. Because we saw enhanced biofilm at either calcium level, we present data from both calcium conditions in this paper and note the calcium level in the figure legends.

**Figure 3.**
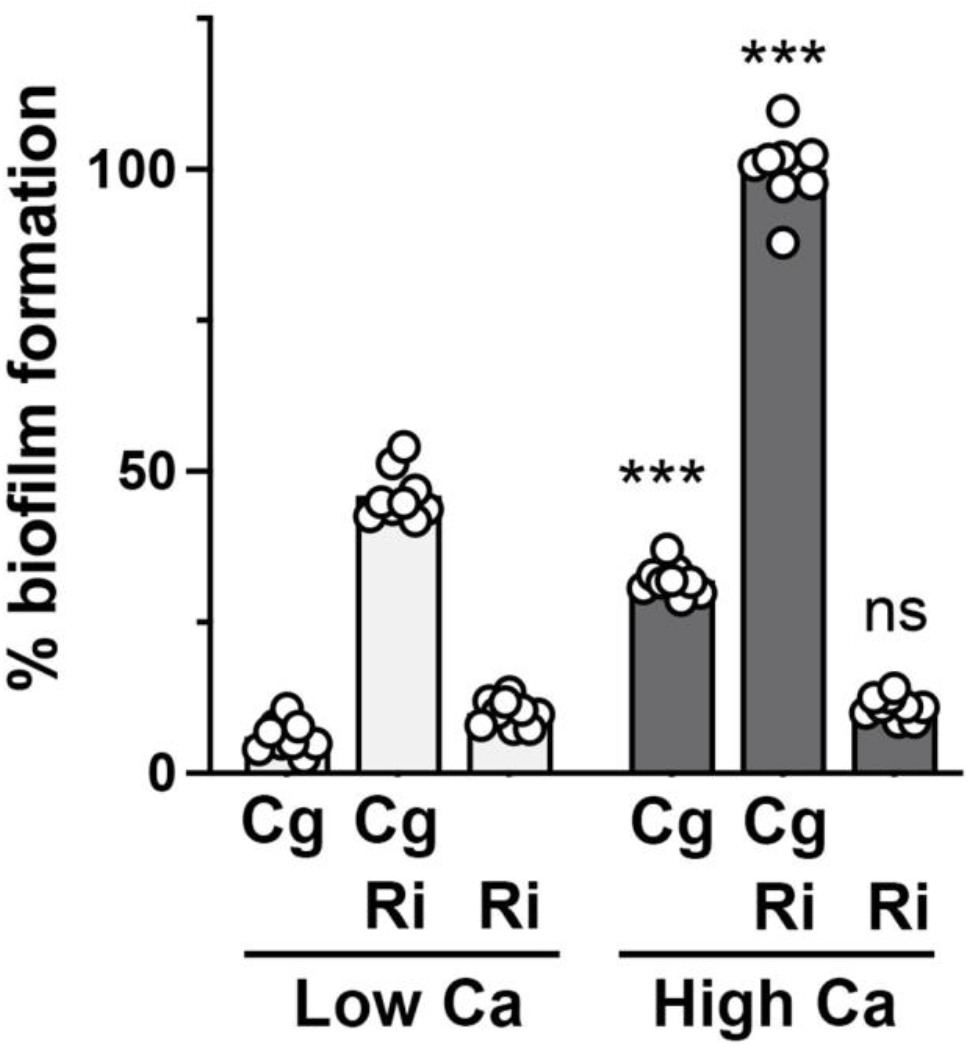
Impact of calcium on biofilm formation by *C. gleum*. More biofilm is generated by *C. gleum* and the dual species biofilm when grown in high calcium (400 μM) MOPS media versus low calcium (21.1 μM) MOPS media. Calcium concentration did not impact *R. insidiosa* biofilm formation. Dots represent individual experimental replicates from three independent experiments; statistic calculated using the means of each experiment. Two-way ANOVA with a Sidak’s multiple comparison test separately comparing each species composition between the two calcium conditions. ***, p < 0.001; ns, not significant.

### The biofilm stimulation between these strains is robust to starting inoculum

Our initial studies used equal starting OD’s for each species and we chose to next examine the robustness of the enhanced biofilm formation to starting inoculum. If we changed the ratio of the starting inoculum of *C. gleum* to *R. insidiosa* from 10% *C. gleum* (90% *R. insidiosa*) to 90% *C. gleum* (10% *R. insidiosa*), where 50% *C. gleum* is our previously reported 1:1 mixture, we saw that all combinations resulted in enhanced biofilm formation compared to *C. gleum* alone (100% *C. gleum*), particularly at the lower proportions of *C. gleum* (**Figure 4A**). Further dilution of *C. gleum* compared to a set proportion of *R. insidiosa* showed that some biofilm enhancement of the co-culture compared to that of *R. insidiosa* alone was retained down to 0.0001% *C. gleum* (roughly 15 *C. gleum* cells per well at time of inoculation) (**Figure 4B**).

**Figure 4.**
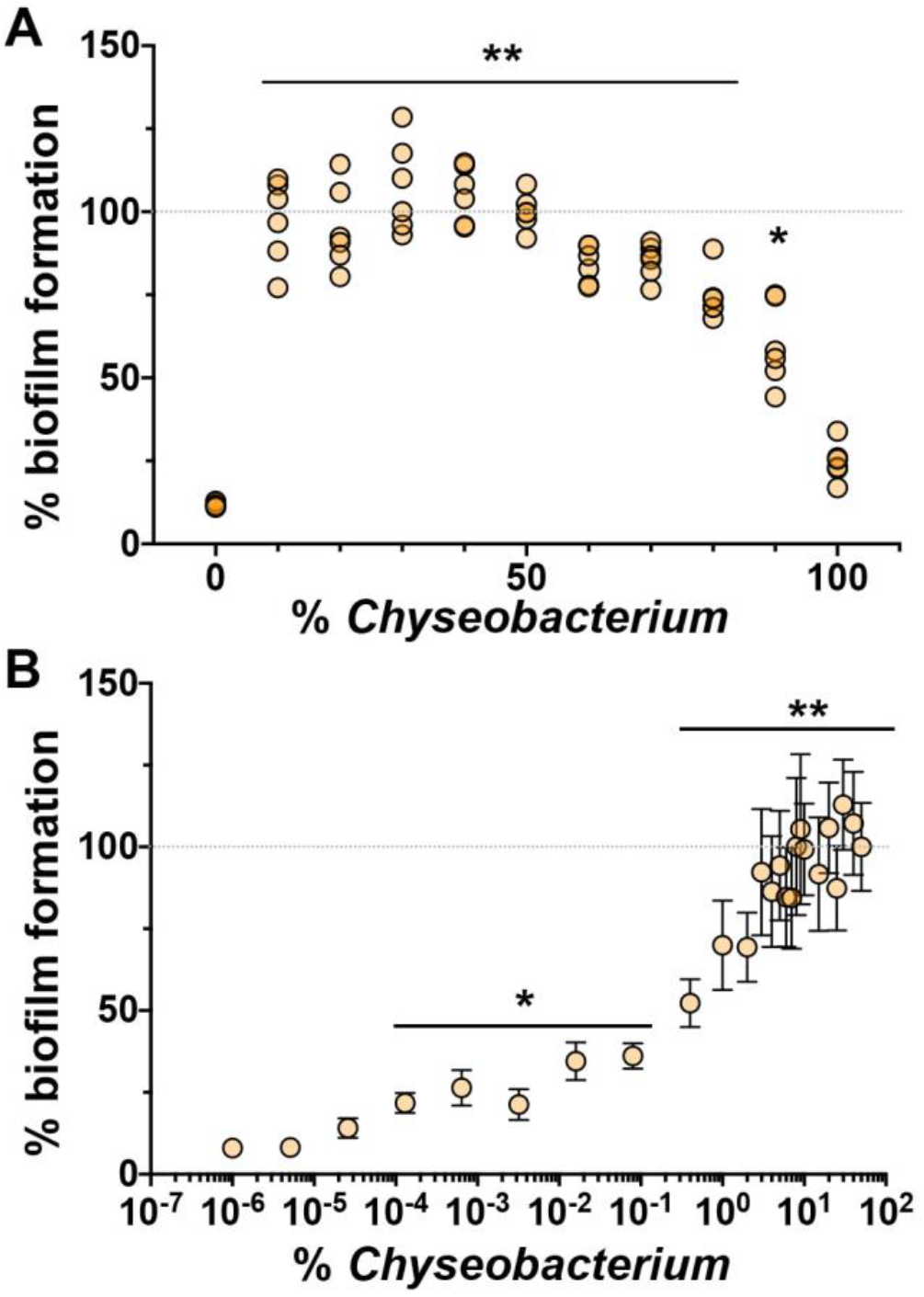
Biofilm stimulation with varying initial concentrations of *C. gleum*. The ISS *C. gleum* was added as a percentage of the initial starting OD_600_ noted on the x-axis with the remainder being comprised of ISS *R. insidiosa*. All experiments conducted in high calcium MOPS **(A)** *C. gleum* stimulation of dual-species biofilm formation decreases as the *C. gleum* composition increases above 50% but shows stable stimulation as *C. gleum* composition drops below 50%. Each dot represents the measurement of a biological replicate from three separate experiments (two replicates per experiment). **(B)** Dilution to extinction of *C. gleum* shows biofilm stimulation above *R. insidiosa* alone (the zero *C. gleum* condition is set as the 10^-6^ dot), showing that biofilm stimulation can be seen with as little as 0.0001% *C. gleum*. Dots represent the mean of at least three independent experiments and the error bars represent standard deviation. The grey dotted line marks the 100% biofilm formation that is set based on the 50/50 *R. insidiosa*, *C. gleum* mix and all other biofilm formation is normalized to this 100% mark. Data were analyzed with Browne-Forsythe and Welch ANOVA with a Dunnett’s post-test with 100% *C. gleum* as the comparator in (A) and 100% *R. insidiosa* as the comparator in (B). *, p < 0.05; **, p <0.01.

### Media and environmental impacts on the *C. gleum/R. insidiosa* interaction

After observing that the enhanced biofilm formation in the *C. gleum/R. indiosa* co-culture was very robust to the starting inoculum (**Fig. 4**) and that calcium concentration could alter *C. gleum* mono- and co-culture biofilm formation (**Fig. 3**), we expanded the parameters and examined the impacts of media composition, temperature, and low-shear simulated microgravity conditions.

In an initial attempt to simplify the media composition, replacement of the MOPS media in our standard MRLP with M9 (M9-RLP) or M63 (M63-RLP) was attempted, as was the RL composition (see Materials & Methods for precise descriptions). When overnight cultures were pre-grown in MRLP and used to inoculate biofilm formation in various media, all tested media supported some level of enhanced biofilm between *R. insidiosa* and *C. gleum* (**Figure 5A**). However, when overnights were pre-grown in M9-RLP, biofilm enhancement was only seen in MRLP and RL media, while no significant biofilm enhancement was seen with M9-RLP or M63-RLP (**Figure 5B**). In data not shown here, pre-growth in RL media was similar to MRLP, while pre-growth in M63-MRLP was similar to M9-RLP. One of the primary differences between MRLP/RL vs M9-RLP/M63-RLP is phosphate concentration, but we have not formally tested if phosphate is the contributing factor in these media differences.

**Figure 5.**
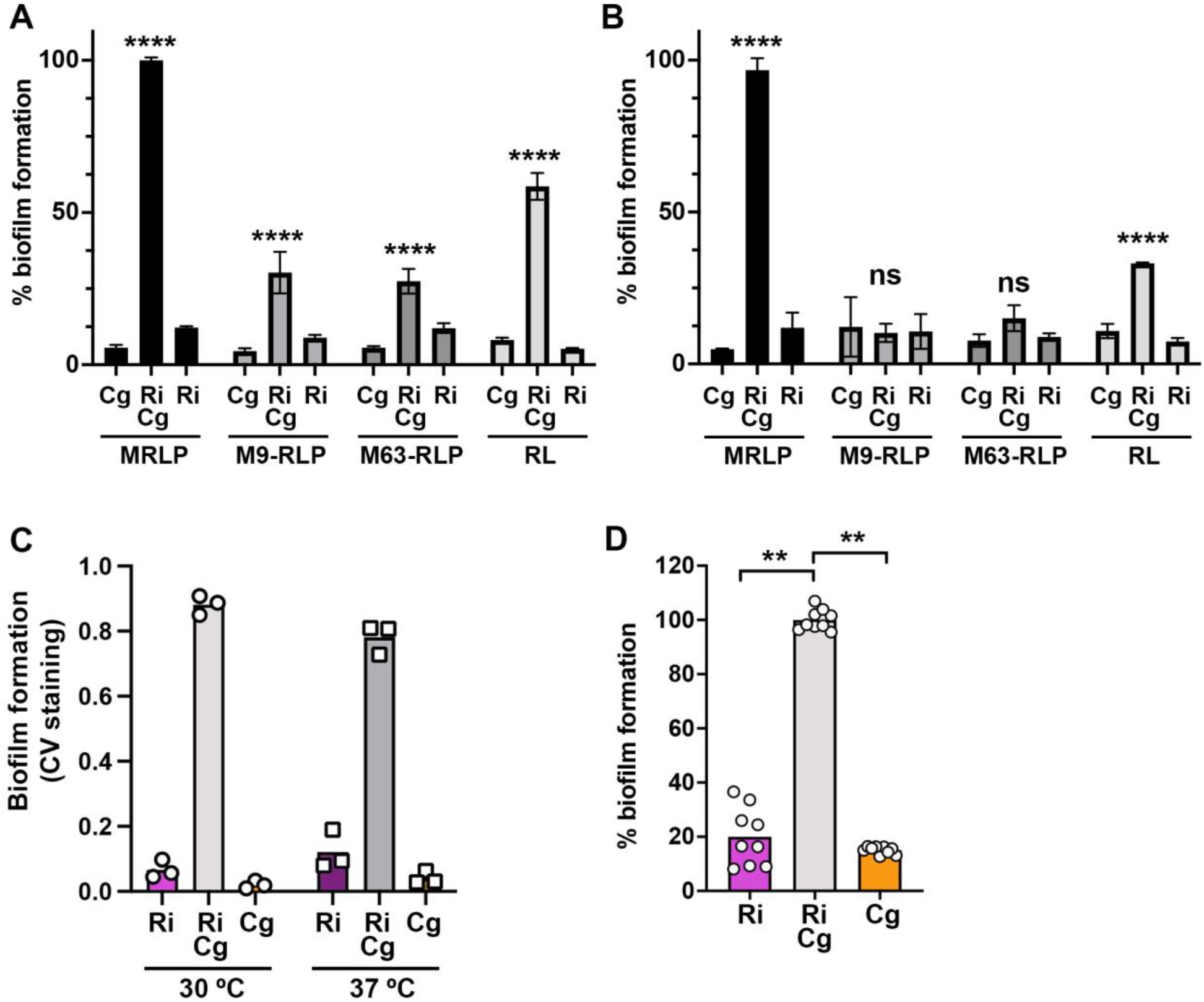
Characterization of growth media, temperature, and low-shear growth on enhanced biofilm formation. **(A)** Biofilm formation normalized to Ri+Cg MRLP for the media listed below the x-axis, with cells pre-grown in low calcium MRLP. **(B)** Normalized biofilm formation in the media listed below the x-axis, with cells pre-grown in low calcium M9-RLP. Data in (B) is normalized to Ri+Cg in MRLP from (A). Means from three independent experimental days, error bars represent standard deviation. **(C)** Biofilm formation from the noted single or dual-species cultures in low calcium MRLP grown at 30°C (left), or 37°C (right). Each point represents the mean of three biological replicates for each of three independent experimental days. **(D)** Biofilm formation on the gas-permeable membrane of a rotating wall vessel (RWV) in high calcium MRLP. Each point represents each replicate of the RWV experiment, though statistics calculated from each experimental day’s mean (i.e. n=3). Statistics for (A,B,C) done using 2-way ANOVA with a Dunnett’s corrected multiple comparison post-test comparing each single species biofilm to the dual-species biofilm within each media or condition. Statistic for (D) using Welch’s ANOVA with Dunnett’s post-test comparing each single species biofilm to the dual-species biofilm.

We also tested whether enhanced biofilm formation by *C. gleum* and *R. insidiosa* was achieved at higher temperatures. Both species are opportunistic pathogens and thus capable of growth at body temperature. When grown at 37°C, similar biofilm enhancement between these species was observed as when grown at 30°C (**Figure 5C**).

Finally, the focus of these studies were two strains collected from the ISS and thus previously subjected to continuous microgravity and the low-shear environment that accompanies such gravity conditions. To mimic the low-shear component of microgravity, we used rotating wall vessels (RWVs) where the cells in suspension have defined vertical circular paths within a low-shear environment. While some clumping was seen for the cells in suspension in the dual-species RWV cultures, we noted that most of the biomass was present as biofilm on the gas-permeable membrane of the RWV chamber. We removed the biofilm from the membranes with silicon cell-scrapers and crystal violet stained the collected cells to assess total membrane attached biofilms. Under these RWV conditions, enhanced biofilm formation was readily apparent compared to the individual species (**Figure 5D**). Note that in the RWV system, *R. insidiosa* forms a more robust biofilm than in most other conditions reported in this study, reminiscent of its stronger biofilm on glass (**Figure 2A**).

### Testing potential mechanisms of biofilm enhancement

Most of the previous descriptions of *R. insidiosa* stimulation of biofilm in mixed cultures showed that the effect was contact dependent and not co-aggregation based. Our microscopy supported comingling of the two species (**Figure 2B**), but we also wanted to formally test whether a soluble factor or co-aggregation could explain the enhanced biofilm formation of this specific interaction. To test whether biofilm stimulation was transferrable by conditioned media, we used cell-free supernatants from *C. gleum, R. insidiosa*, or co-cultures mixed 1:1 with fresh media and measured biofilm formation in the flexible 96-well format. Neither single species nor the dual-species supernatants were able to stimulate biofilm formation (**Figure 6A**). This supports the direct interaction model but does not fully rule out potential for a soluble mediator, especially if the mediator is short-lived.

**Figure 6.**
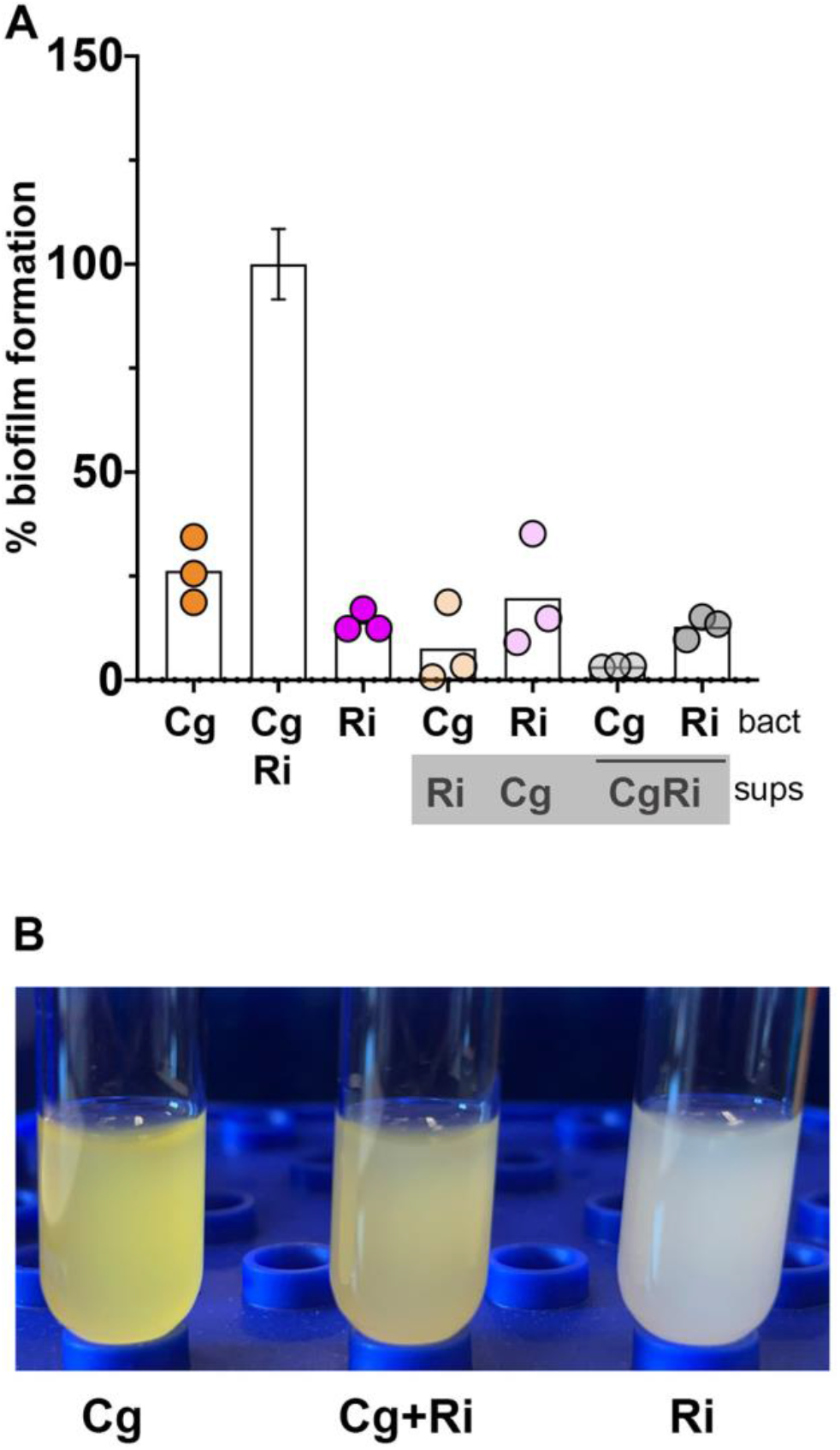
The biofilm enhancing effect of either species cannot be transferred by supernatant and is not due to co-aggregation. **(A)** The enhancement in biofilm formation seen in the dual-species growth is not conferred by cell-free supernatant from either the single or dual-species conditions using 50% spent supernatant and 50% fresh media. None of the supernatant treatments were significantly different than their single species growth after correction for multiple comparisons using a Brown-Forsythe and Welch ANOVA with a Dunnett’s post-test comparing all means to each other. The source of the supernatants tested is present in the grey box under the species abbreviation. Each dot represents the mean from three biological replicates per experiment. The Cg/Ri conditions is set to 100% for each experiment to normalize between experiments, so the average standard deviation for 50/50 Cg-Ri mixes is displayed. **(B)** There is no co-aggregation apparent between these strains. Cells grown in the high-Ca MOPS media were collected by centrifugation and resuspended at a final concentration of OD_600_ = 6 and examined over time with the picture representative of all replicates at one hour post mixing.

We assessed the potential for co-aggregation by mixing concentrated cultures of both species together and looking for any sedimentation caused by aggregation. At one hour there was no evidence of sedimentation (**Figure 6B**) and no sedimentation indicative of aggregation was apparent at times up to 24 h (data not shown).

### Species and strain specificity for dual-species biofilm enhancement

Our primary goal was to characterize the interaction between the ISS *C. gleum* and *R. insidiosa* to understand our potable water model community. However, we also wanted to understand how broadly applicable this interaction was for other species in these genera and other strains of these species. We acquired non-ISS strains of *R. insidiosa, R. pickettii*, an unnamed *R. sp., C. gleum, C. indologenes*, and *C. meningisepticum* (**Table 1**) and conducted biofilm assays in the flexible 96-well dish format. Most of the *R. insidiosa* strains could enhance ISS *C. gleum* biofilm formation, while only some of the other two Ralstonia species were able to do this (**Figure 7A, B**). The ability of ISS *C. gleum* to respond to Ralstonia presence with enhanced biofilm formation, however, was limited only to ISS *C. gleum* and very mild stimulation of *C. meningisepticum* biofilm (**Figure 7A, C**). A big difference between the ISS *C. gleum* and the other strains is that the other Chryseobacterium tested, particularly the *C. gleum* strains, form robust biofilms on their own (**Figure 7A**).

**Figure 7.**
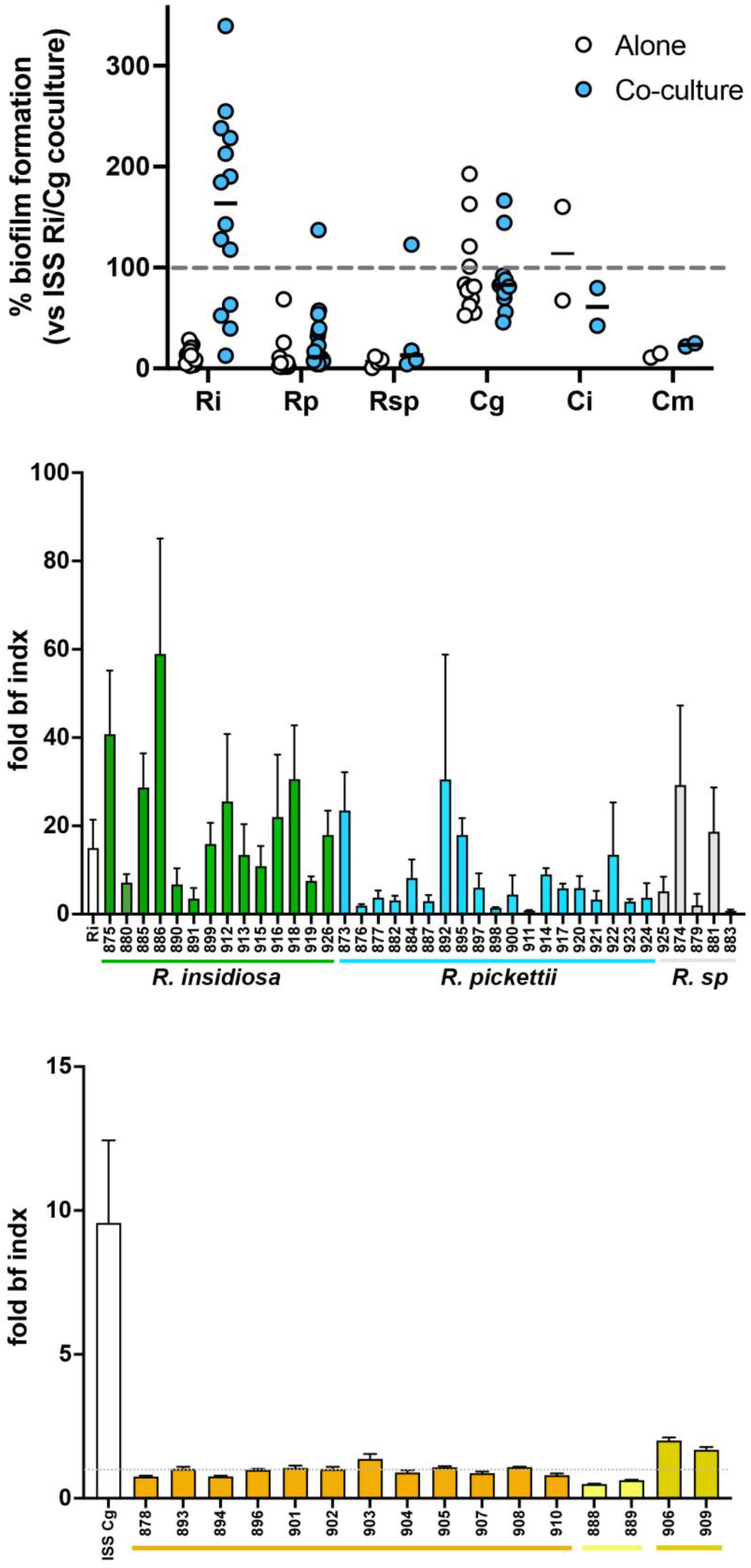
Strain and species specificity for dual-species biofilm enhancement. **(A)** Biofilm quantification normalized by setting the intraday ISS *R. insidiosa* and ISS *C. gleum* dual-species biofilm as 100% biofilm formation. White circles are data for each species alone and the blue circles are in the presence of either the ISS *C. gleum* (for Ralstonia species) or the ISS *R. insidiosa* (for the Chryseobacterium species). Two-way repeated measures ANOVA with Sidak’s multiple comparison test comparing intra-species mono- and dual-species biofilms supports that at the species level, only *R. insidiosa* shows significant biofilm enhancement under dual species conditions (p < 0.001). **(B)** The same data contributing to panel (A) converted to fold change of dual species biofilm divided by biofilm formation by that strain alone. Green bars are the *R. insidiosa* strains, blue are *R. pickettii* strains, and grey are an unnamed Ralstonia species. **(C)** The same data contributing to panel (A) represented as fold change of dual species biofilm divided by biofilm formation by that strain alone. The white bar is the ISS *C. gleum*, orange bars are the non-ISS *C. gleum* strains, yellow are *C. indologenes* strains, and mustard are *C. meningisepticum* strains.

## Discussion

Here we examined the interaction between ISS-derived *R. insidiosa* and *C. gleum* to better understand the enhanced biofilm of the dual-species cultures (**Figure 1**) as part of community biofilm formation in our six-species potable water community [24]. The enhanced biofilm formation between these two ISS strains is robust to starting inoculum (**Figure 4**) and temperature (**Figure 5C**), does not occur in all growth media (**Figure 5B**), and does not appear to be driven by a soluble mediator or co-aggregation (**Figure 6**). While these findings shed light on the ISS *R. insidiosa* and *C. gleum* interaction, such enhancement is not a common interaction outcome between these species based on our examination of other *R. insidiosa* and *C. gleum* strains, as well as other species of Ralstonia and Chryseobacterium (**Figure 7**). Thus, while the work presented here increases our understanding of the potable water model system, we fully acknowledge that most of the findings are not broadly extrapolatable to strains found outside of the ISS.

The enhanced biofilm formation between the ISS *R. insidiosa* and *C. gleum* is readily apparent (**Figure 1A**) and is not driven by changes in total cell number or proportional changes in cell number in the cultures (**Figure 1B**). The material presented here is part of a broader characterization of the ISS potable water model community, and thus has been underway for quite a number of years and with many different graduate, undergraduate, and technician contributors. It was during data examination for this report that we realized that there were two different MOPS media recipes co-existing in the lab differing only by their calcium concentrations. This calcium difference was sufficient to describe the higher biofilm formation by *C. gleum* monoculture observed by some lab members but not others (**Figure 3**). The general biofilm enhancement between the two ISS bacteria is still readily apparent at either calcium concentration, and the calcium concentration of the resultant MRLP media in each experiment is noted in each figure legend.

The enhancement of biofilm formation between the ISS *R. insidiosa* and *C. gleum* is very robust to starting inoculation (**Figure 4**). While we did not assess resulting CFUs for each species in these broader inoculation tests, we have no *a priori* reason to suspect that there are different mechanisms of enhancement at different starting inocula, though we cannot rule that out. The enhanced biofilm formation was seen at both 30°C and 37°C (**Figure 5C**) and occurred at room temperature (22-24°C) in the RWV experiments (**Figure 5D**). The RWVs were used to ensure that the biofilm enhancement we observed in open, static biofilms was retained in a system mimicking the low shear environment found during growth in microgravity, as applicable to the ISS. While enhanced biofilm formation could happen with surprisingly low inoculum of *C. gleum*, no enhancement of biofilm could be seen in either direction by single culture or co-culture supernatants (**Figure 6A**), suggesting that there is either no soluble mediator or that any such mediator is very labile. Co-aggregation, while a common mechanism in biofilms particularly in the oral cavity, does not drive the enhanced biofilm formation in this interaction (**Figure 6B**).

When the ISS strains were grown in MRLP in the overnight cultures used for inoculation, there was observable dual-species biofilm enhancement in all tested media (**Figure 5A**), but when cells were instead grown in overnights of M9-RLP (**Figure 5B**) or M63-RLP (not shown) there was no biofilm enhancement in the subsequent M9-RLP or M63-RLP conditions, though enhancement was retained in MRLP and RL. We suspect that this is due to the relatively high phosphate concentration in the M9 and M63 media that could be suppressing biofilm enhancement. However, mechanistic dissection of the media component contribution to biofilm enhancement was halted once we determined that this interaction was not commonly shared between strains of these species (**Figure 7**).

The ISS *R. insidiosa* behaves similarly to most of the other *R. insidiosa* strains in terms of their weak monoculture biofilm formation and ability to enhance biofilm when grown with ISS *C. gleum* (**Figure 7**). Most of the *R. pickettii* strains do not show strong biofilm enhancement with *C. gleum*, nor does a *Ralstonia pickettii-like* species that has not yet been named. These similarities amongst the *R. insidiosa* strains suggest the ISS *R. insidiosa* and our mScarlet labeled derivative are potentially good representatives of the species. It is at least minimally genetically tractable, as we were able to generate an *attTn7:: mScarlet* integrant that may be useful for studying *R. insidiosa* interaction studies with other bacteria.

The non-ISS *C. gleum* strains form strong biofilms, as shown in **Figure 7** and from literature reports [17], as do the two tested *C. indologenes* strains, while the two tested *C meningisepticum* strains do not. Thus, the ISS *C. gleum* appears to be non-representative of other *C. gleum* strains. While all our Chryseobacterium isolates tested were from clinical sources, others in the literature were isolated from environmental sources and also show strong biofilm formation in monoculture. Thus, our ISS *C. gleum* appears only useful to describe its specific interactions amongst the ISS potable water bacteria.

In conclusion, we have presented partial dissection of an important interaction in a model potable water community. Despite the observations that this strain-specific interaction is unlikely to be broadly applicable beyond this model, we have provided additional information on basal biofilm formation amongst diverse *C. gleum* and *R. insidiosa* strains as well as generated an mScarlet-expressing *R. insidiosa* that may be useful to others in the field or studying *R. insidiosa* pathogenesis.

## Acknowledgements

We’d like to acknowledge the many undergraduate researchers and graduate rotation students who examined various aspects of the *R. insidiosa/C. gleum* interaction that are not reported here but still contributed holistically to our understanding of these two bacteria, including: Matthew Bompastore, Sierra Bruno, Brynn Cairns, Matthew Kinahan, Alexis Nadeau, Hannah Schulman, Alex Thompson, Sophie Unger, and Trevor Wolf.

